# Low concentrations of tetrasodium EDTA cause significant killing of biofilm-associated *P. aeruginosa* in high validity models of chronic wound and CF lung infections – but not in a model of endotracheal tube colonisation

**DOI:** 10.1101/2025.11.08.687333

**Authors:** Oluwatosin Qawiyy Orababa, Charlotte Cornbill, Ayomikun Kade, Natasha Reddy, Rupika Gulati, Freya Harrison

## Abstract

*Pseudomonas aeruginosa* is a pathogen notorious for its antimicrobial resistance and is currently classified as a high-priority pathogen for which new drugs are needed. Tetrasodium EDTA (tEDTA) is one of the new antimicrobial compounds that have been shown to have good antibacterial and antibiofilm efficacy against *P. aeruginosa*. Due to the diversity and highly drug-tolerant nature of *P. aeruginosa* biofilms in different infection environments, it is important to carry out pre-clinical testing of new antibiofilm agents against this pathogen in media and models that accurately mimic diverse infection environments. In this study, we used different high validity media and biofilm models that mimic chronic wounds, endotracheal tubes, and cystic fibrosis lung infections to assess the efficacy of tEDTA against *P. aeruginosa* biofilms. We report that different infection environments influence the susceptibility of both planktonic and biofilm forms of *P. aeruginosa* to tEDTA. The highest tolerance to tEDTA was observed in the media and biofilm model that mimics the endotracheal tube environment. In conclusion, we show that although different infection environments influence the efficacy of tEDTA against *P. aeruginosa* biofilms, it has good potential for use as an alternative antimicrobial in treating *P. aeruginosa*-associated biofilm infections.

## INTRODUCTION

Antimicrobial resistance (AMR) remains one of the greatest challenges of the 21st century with increasing mortality and economic cost. In 2021 alone, there were about 4.71 million deaths associated with bacterial AMR, and this figure has been predicted to rise and reach around 10 million annual deaths by 2050 (Naghavi et al., 2024). Additionally, bacterial AMR accounted for a total hospital cost of US$693 billion and a productivity loss of US$194 billion globally in 2019 (Naylor et al., 2025). As a result, there is a need for novel classes of antimicrobials and alternative therapeutic options to effectively treat AMR/highly tolerant infections, especially those associated with Gram-negative bacteria (Prasad et al., 2022). Typical examples of AMR infections are those associated with biofilms. Biofilm-associated infections account for about 65% of all bacterial infections and 80% of all chronic, non-healing microbial infections (Jamal et al., 2018). These infections, including chronic wounds, ventilator-associated pneumonia, and cystic fibrosis lung infections, are highly tolerant to antibiotics due to the presence of the biofilm matrix and physiological changes associated with the biofilm state (Shree et al., 2023).

*Pseudomonas aeruginosa* belongs to the ESKAPE (*Enterococcus faecium, Staphylococcus aureus, Klebsiella pneumoniae, Acinetobacter baumannii, P. aeruginosa*, and *Enterobacter* spp) pathogen list of bacteria that are highly resistant to antibiotics (Miller et al., 2024). It is also a high-priority pathogen for which new treatments need to be developed, and forms highly robust biofilms in sites of infection. *P. aeruginosa* forms biofilms using three main exopolysaccharides (alginate, Pel, and Psl), and this may vary depending on the infection site and strain (Gheorghita et al., 2023; Chung et al., 2023). Alginate-encased biofilms are often associated with CF lung infections, while Pel plays a more significant role in wound infections (Chung et al., 2023). Consequently, susceptibility of *P. aeruginosa* biofilms to antibiotics is predicted to be influenced by the infection environment and biofilm matrix type.

Previous research has clearly demonstrated the influence of infection environments on bacterial physiology and their susceptibility to antimicrobials (Ersoy et al., 2017; Kubicek-Sutherland et al., 2015; van den Bossche et al., 2021; Hasan et al., 2022). Previous research by our team on in-use antibiotics and candidate novel antibiofilm agents against *P. aeruginosa* has shown discordance between assays conducted in rich medium vs. infection-mimicking medium, and between polystyrene plate-based assays using infection-mimicking medium and high-validity *in vitro* and *ex vivo* infection models (Garcia-Maset et al., 2025; Sweeney et al., 2020). It is therefore important to use the right testing models during pre-clinical assessment of antimicrobials with potential to be used in alternative therapies for specific biofilm infection contexts.

Tetrasodium EDTA (tEDTA) has been previously shown to have good antibacterial and antibiofilm activity against various pathogens, including *P. aeruginosa* (Liu et al., 2018; Crowther et al., 2025). It has also been suggested to be a good treatment for wound care (Finnegan and Percival, 2015). However, previous studies of tEDTA have been conducted in media and biofilm models that do not recapitulate *in vivo* infection environments. Liu et al. and Crowther et al. grew biofilms attached to polystyrene pegs or microtitre plates, respectively, in rich laboratory growth media.

In this study, we assessed the antibacterial and antibiofilm efficacy of tEDTA against *P. aeruginosa* in three high-validity models of specific biofilm infection contexts, which are in regular use in our lab. These were an *in vitro* model of soft-tissue chronic wound biofilm (Werthén et al., 2010), a porcine *ex vivo* model of biofilm in the lungs of people with cystic fibrosis (Harrington et al., 2021) and an *in vitro* model of biofilm in the endotracheal tubes (ETTs) used to connect hospital patients to ventilators (Walsh et al., 2024a). We used a well-characterised laboratory strains of *P. aeruginosa*: PA14, which can produce the pel and alginate polysaccharides as well as proteinaceous and nucleic acid matrix components. We showed that although tEDTA has good antibiofilm efficacy against *P. aeruginosa* in the different models, the level of efficacy varies by strain and growth environment. Biofilms grown in the ETT model were notably harder to eradicate with tEDTA than biofilms grown in the other two models.

## RESULTS AND DISCUSSION

### Tetrasodium EDTA has good antibacterial efficacy against planktonic *P. aeruginosa* in standard rich medium and host-mimicking media

Environmental cues in various infection environments play crucial roles in determining bacterial susceptibility to antibiotics (Ersoy et al., 2017; Kubicek-Sutherland et al., 2015; van den Bossche et al., 2021; Hasan et al., 2022; Garcia-Maset et al., 2025; Sweeney et al., 2020). To determine the impact of growth medium on the susceptibility of *P. aeruginosa* to tEDTA, we first assessed its minimum inhibitory concentration (MIC) against planktonically-grown *P. aeruginosa* PA14 using the broth microdilution assay in both standard laboratory media (caMHB) and infection-mimicking media. The infection-mimicking media were simulated wound fluid (SWF, Werthén et al., 2010), synthetic cystic fibrosis sputum media (SCFM1, Palmer et al, 2007), and synthetic ventilated airway mucus (SVAM, Walsh et al. 2024b) – these media can be used to make high-validity biofilm platforms that mimic the environments of soft-tissue wounds, CF airways and endotracheal tubes, respectively.

tEDTA inhibited the growth of both *P. aeruginosa* strains in all the media tested with MICs ranging from 0.25% to 1% (Table 1). The highest MIC was seen with PA14 in the SVAM; this strain had a four-fold variation in MIC across the media tested. This evidenced the role of growth environment in the susceptibility of this pathogen to tEDTA, even though the variation between media tested was relatively small. The antibacterial efficacy of tEDTA against *P. aeruginosa* in this study is in line with previous studies that have reported that this compound has good efficacy against *P. aeruginosa* (Sivaranjani et al., 2021; Crowther et al., 2025). The MIC (0.25%) reported in this study in both the standard lab medium (caMHB) and SWF is also similar to the MIC reported by Sivaranjani et al. (2021). However, we report a slightly higher (0.5-1%) MIC in SCFM and SVAM against *P. aeruginosa* PA14.

**Table 1.** Minimum inhibitory concentrations of tetrasodium EDTA in different media.

|  | caMHB | SWF | SCFM | SVAM |
| --- | --- | --- | --- | --- |
| PA14 MIC (% w/v) | 0.25 | 0.25 | 0.5 | 1 |

### Tetrasodium EDTA causes significant killing of mature *P. aeruginosa* biofilm, in a soft tissue chronic wound model

*P. aeruginos*a is one of the major bacterial pathogens in chronic wound infections and is associated with treatment difficulty (Rahim et al., 2017). We assessed the ability of tEDTA to inhibit biofilm formation or to eradicate an established biofilm of *P. aeruginosa* PA14 strain in a soft tissue chronic wound model (Fig 1A) (Werthén et al., 2010). Werthén et al. (2010) previously showed that the structure of *P. aeruginosa* biofilm formed in this *in vitro* model is similar to its structure *in vivo*. We also allowed biofilms to grown in the model for 24h, before adding the treatment for a further 24h. Both 0.5% tEDTA and 1% tEDTA caused a more than 3-log_10_ reduction in biofilm population (Fig 1B). This is the first report of the biofilm eradication efficacy of tEDTA in this chronic wound model. Crowther et al. (2025) showed that treatment of *P. aeruginosa* biofilm with 4% tEDTA resulted in a 3-log_10_ reduction in biofilm population using the polystyrene microtitre plate platform and rich growth medium. However, in our study, we observed 3-log_10_ reduction with a much lower tEDTA concentration than that used by Crowther et al. (2025) – indicating lower tolerance of biofilm to tEDTA in the wound model. Also, the variations in concentration might be linked to differences in *P. aeruginosa* strain used. Crowther et al. (2025) used PAO1, while we used the PA14 strain in this study. These two strains produce different biofilm-associated exopolysaccharides (Gheorghita et al., 2023; Chung et al., 2023).

**Fig 1.**
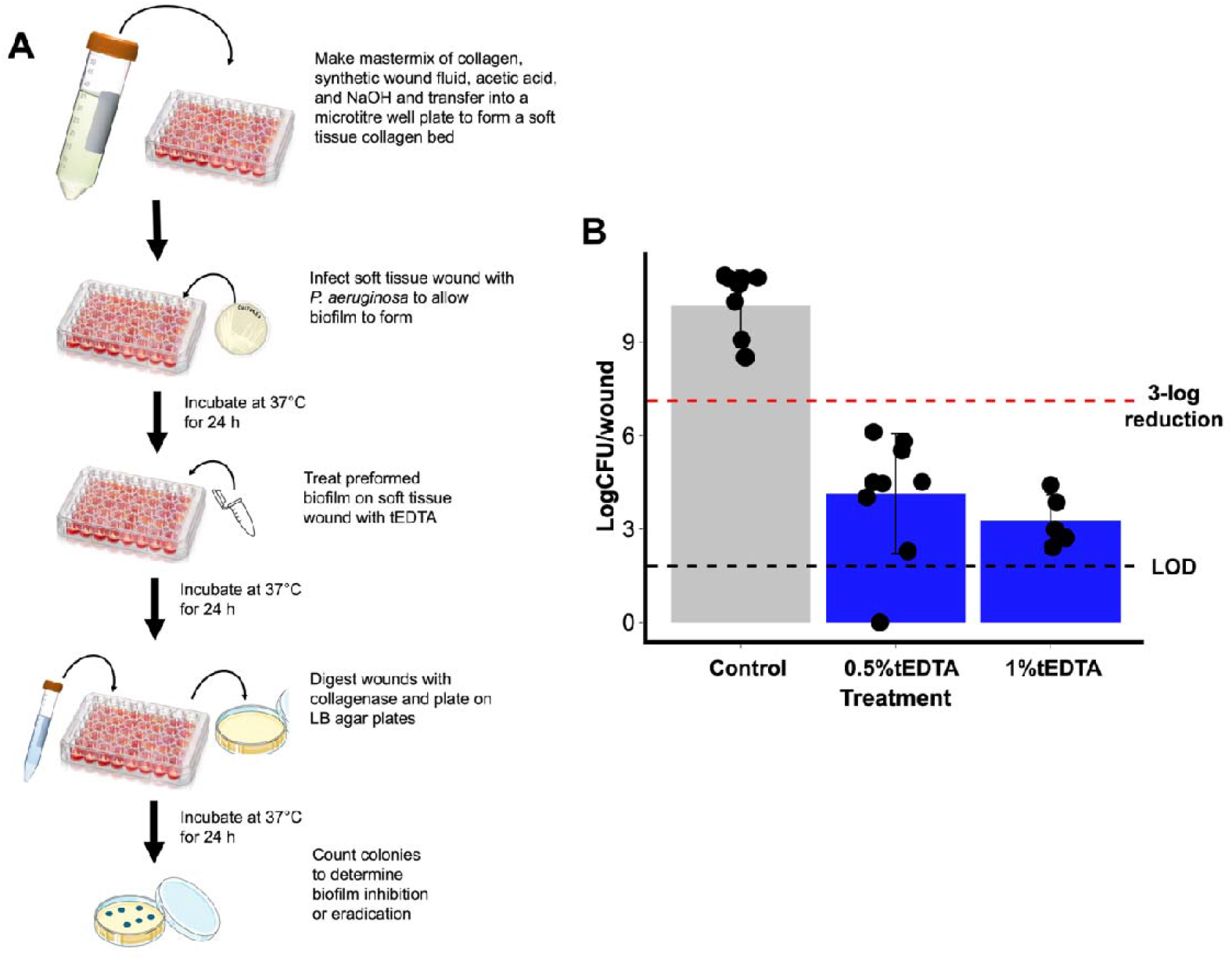
Assays for prevention and eradication of *P. aeruginosa* biofilm in a soft tissue chronic wound model. **A**. Flow diagram showing how the soft tissue wound biofilm model was used for assessing biofilm inhibition and eradication. **B**. A soft tissue wound was infected with *P. aeruginosa* PA14 and incubated for 24 h to allow biofilms to form. The preformed biofilms were treated with different concentrations (0.5% and 1%) of tEDTA and incubated at 37°C for another 24 h. Bacteria were recovered by enzymatic digestion of the collagen and colony-forming units (CFU) counted to determine viable cell numbers. The red dashed line indicates 3-log_10_ bacterial reduction; the black line indicates the limit of detection (LOD) for CFU by plating. Error bars are mean ±SD.

### Tetrasodium EDTA can completely eradicate mature *P. aeruginosa* biofilm in a CF lung model

Chronic *P. aeruginosa* lung infection is prevalent among adults with CF, and is responsible for a significant decrease in health and life expectancy (Durda-Masny et al., 2021; Durfey et al., 2025). We assessed the ability of tEDTA to eradicate preformed biofilm of *P. aeruginosa* in a CF lung model (Fig 2A) which has been shown to induce high-level tolerance to antibiotics (Harrington et al., 2020; Harrington et al., 2021; Sweeney et al., 2020). tEDTA caused more than 3-log_10_ reduction in biofilm population of *P. aeruginosa* PA14 when used at 1% in this model (Fig 2B). Interestingly, 2% tEDTA completely eradicated *P. aeruginosa* biofilm in this model (Fig 2B). To our knowledge, no previous study has reported the effect of tEDTA on CF lung pathogens. The good biofilm eradication efficacy we see with tEDTA against *P. aeruginosa* in this CF lung model indicates its potential for development into a nebulised treatment or a sinus wash that could manage CF lung infections resulting from *P. aeruginosa*.

**Fig 2.**
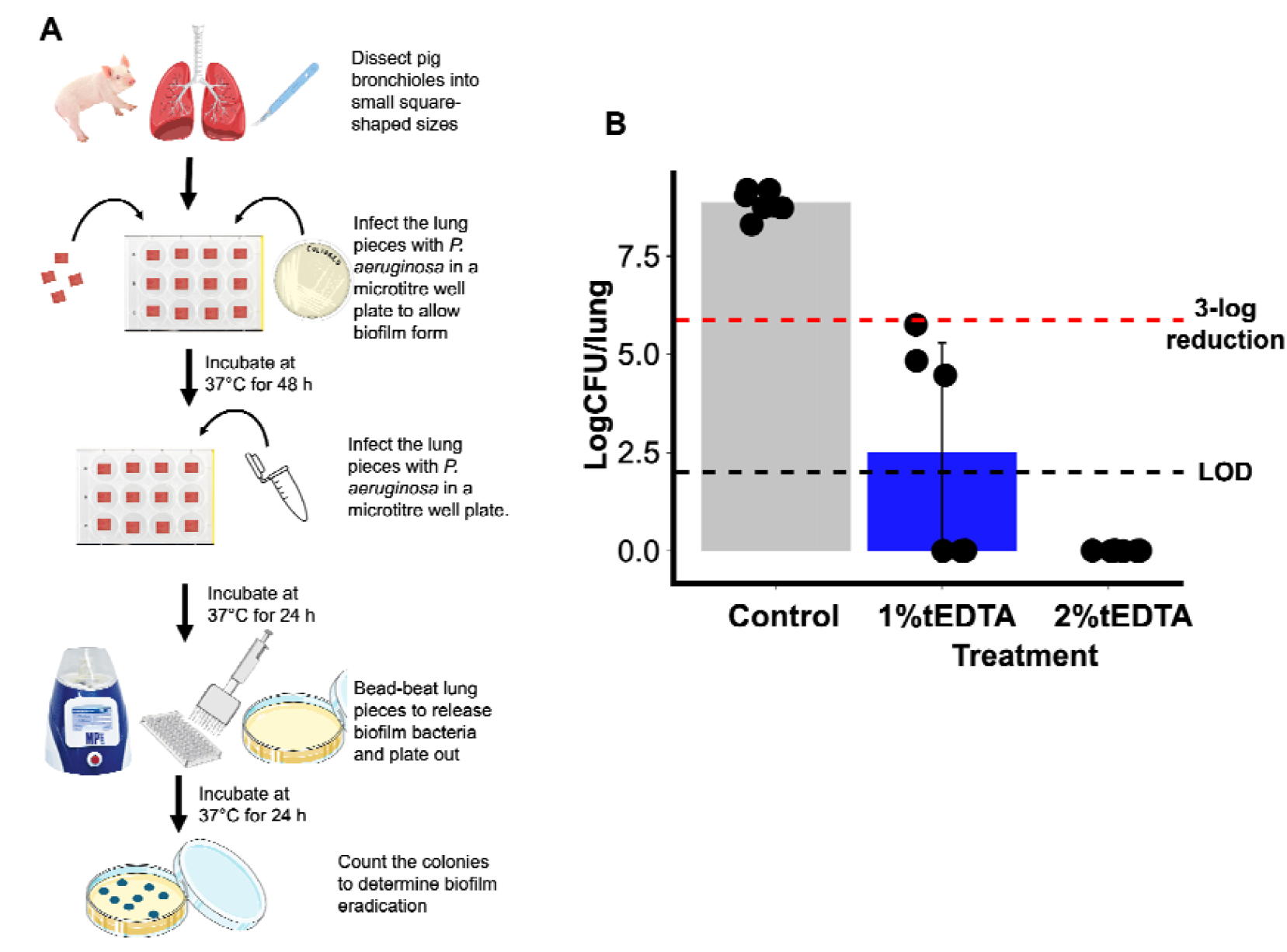
Assay for eradication of *P. aeruginosa* biofilm in an *ex vivo* cystic fibrosis lung model. **A**. Flow diagram showing how the *ex vivo* pig lung (EVPL) model of CF infection was used to assess biofilm eradication. **B**. Bronchioles from pig lungs were dissected into small pieces and infected with *P. aeruginosa* PA14. The infected lung was incubated at 37°C for 48 h to allow biofilms to form. After 48 h, the lungs were treated with different concentrations (1% and 2%) of tEDTA and incubated for a further 24 h before bacteria were recovered from the tissue-associated biofilm by bead beating and colony-forming units (CFU) counted to determine viable cell numbers. The red dashed line indicates 3-log_10_ bacterial reduction; the black line indicates the limit of detection (LOD) for CFU by plating. Error bars are mean ±SD.

### Biofilm eradication of *P. aeruginosa* with tEDTA is more difficult in an endotracheal tube model

Ventilator-associated pneumonia (VAP) is one of the most common nosocomial infections in critically ill patients, with *P. aeruginosa* being a common pathogen in VAP (Howroyd et al., 2024). VAP is ultimately a consequence of biofilm formation on the ETT used to connect the patient to the ventilator: fragments of the ETT-associated biofilm colonise the lungs, leading to pneumonia (Durairaj et al., 2009; Pneumatikos et al., 2009; Diaconu et al., 2018). Using an *in vitro* ETT (IVETT) model that was recently developed in our lab (Walsh et al., 2024a), we assessed the ability of tEDTA (2%, 3%, and 4%) to eradicate mature *P. aeruginosa* biofilm on an abiotic surface (Fig 3A). Among the tested concentrations, only 4% tEDTA yielded more than a 3-log_10_ reduction in viable bacteria (Fig 3B). This was higher than the concentration required to achieve at least 3-log_10_ biofilm killing in the wound (0.5%) or EVPL (1%) models. However, this concentration (4%) is deemed safe and is currently used in the clinic to prevent bloodstream infections associated with central venous catheters (Robinson et al., 2025). The concentration of tEDTA required to achieve a 3-log_10_ reduction in biofilm population in the IVETT is the same as the concentration required by Crowther et al. (2025) to cause a comparable killing of biofilms grown in polystyrene microtitre plates. Although no study has previously checked the antibiofilm efficacy of tEDTA in the IVETT model, previous work has explored the impact of tEDTA on other infections associated with medical device biofilms. For example, Robinson et al. (2025) showed that 4% tEDTA prevents central-venous catheter-associated bloodstream infections in paediatric haemodialysis patients. Moore et al. (2023) also showed that about 1% tEDTA reduced the risk of catheter blockade with biofilms in an *in vitro* catheter-associated urinary tract infection model. Similarly, Hirsch (2024) showed that 4% tEDTA reduced the risk of composite catheter complications in paediatric patients.

**Fig 3.**
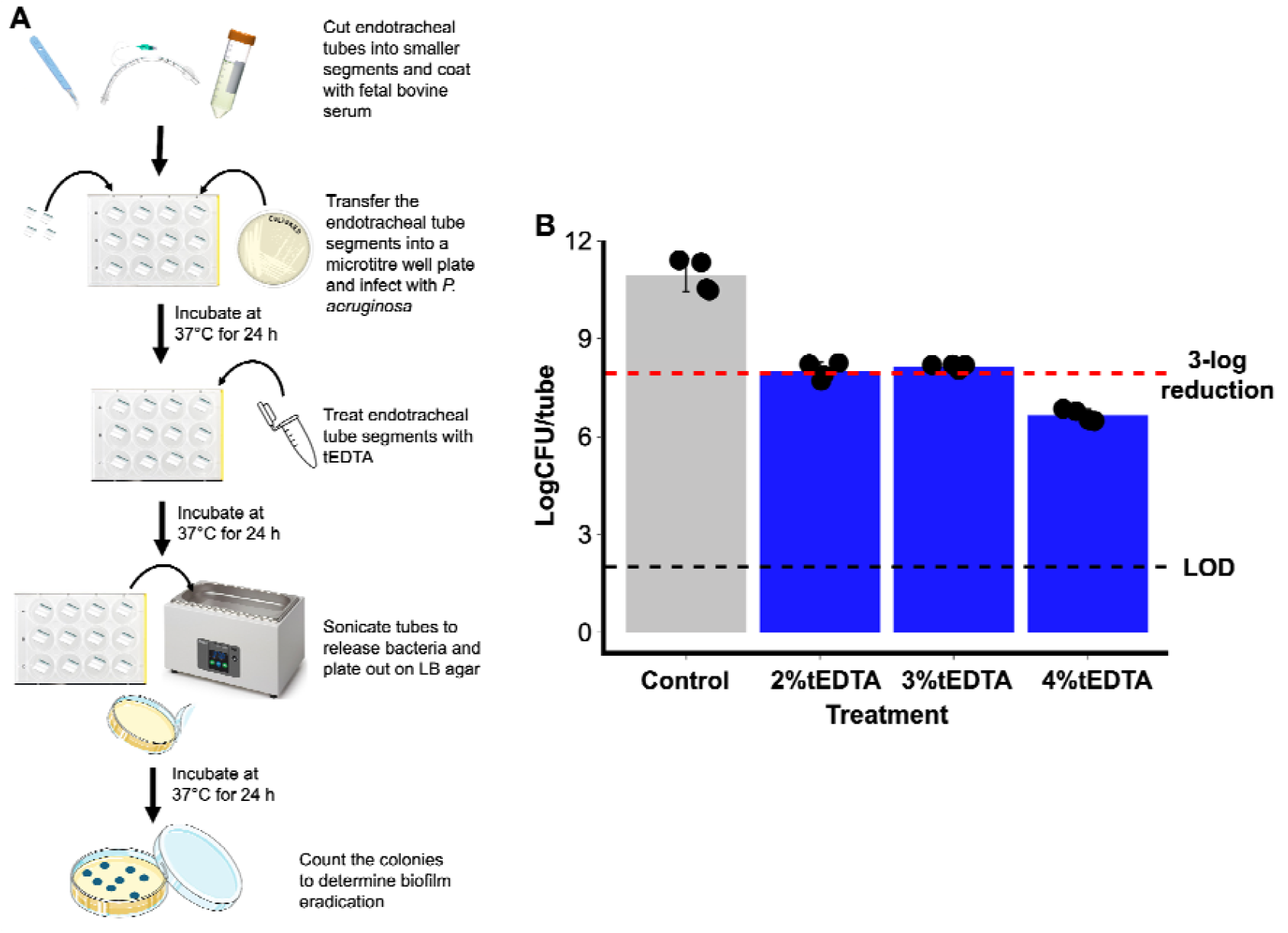
Assay for eradication of *P. aeruginosa* biofilm with tEDTA in an *in vitro* endotracheal tube (IVETT) biofilm model. **A**. Flow diagram showing how the IVETT model was used to assess biofilm eradication. **B**. Segments of endotracheal tubes were coated with fetal bovine serum (FBS), infected with *P. aeruginosa* PA14, and incubated for 48 h in a synthetic ventilated airway mucus (SVAM). Tubes were then treated with specific concentrations (2%, 3%, and 4%) of tEDTA for 24 hours. After this, bacteria were recovered from the ETT-associated biofilm by sonication and CFU enumerated. The red dashed line indicates 3-log_10_ bacterial reduction. Error bars are mean ±SD.

## CONCLUSIONS

In this study, we showed that different biofilm infection environments are likely to influence the susceptibility of *P. aeruginosa* biofilm to tEDTA. Nonetheless, this non-antibiotic antimicrobial has the potential to be used to treat some *P. aeruginosa* biofilm infections. We have also shown that the antibiofilm efficacy of at least 3-log_10_ is achieved at concentrations below the already clinically approved concentration (4%) of tEDTA in catheter-associated bloodstream infections. Future studies should focus on the suitability of tEDTA for incorporation into wound care products, or into solutions that could be nebulised into the CF lung, or used as washes for sinus decontamination in CF. We did not find promising activity when we tried to eradicate ETT biofilms of *P. aeruginosa* with tEDTA at concentrations of up to 4%. Future work could also usefully explore the ability of tEDTA to prevent or slow biofilm colonisation of ETTs; this may be achievable and could potentially translate into development of coated ETTs that are functionalised with tEDTA. Similarly, experiments assessing the ability of lower concentrations of tEDTA may indicate promise for the prevention of *P. aeruginosa* colonisation of wound infections or CF lungs. Parallel safety/toxicity testing with human cell lines or tissues should ideally be conducted. The mechanism by which EDTA affects bacteria/biofilm is thought to be by chelating cations (Ca^2+^ and Mg^2+^) on the membrane, which likely disrupts bacterial membrane potential (Leive, 1965). Future studies should also identify the specific mechanism of action of tEDTA against *P. aeruginosa*. In conclusion, our work shows the potential of tEDTA as an antimicrobial agent for different *P. aeruginosa*-associated biofilm infections and how different infection environments can influence its antibiofilm efficacy against *P. aeruginosa*.

## METHODS

### Minimum inhibitory concentration assay

The broth microdilution method was used to determine the minimum inhibitory concentration (MIC) of tEDTA (Sigma-Aldrich) as recommended by the European Committee on Antimicrobial Susceptibility Testing (EUCAST). Briefly, the bacterial strain of interest was streaked on Luria-Bertani (LB) agar and incubated at 37°C for 18-24 h to produce distinct colonies. Twice the maximum concentration of tEDTA to be used was prepared in the medium of interest (caMHB, SWF (Werthén et al., 2010), SCFM1 (Palmer et al., 2007), and SVAM (Walsh et al., 2024a,b)) and dispensed (100 µl) in the first column of a Corning Costar CLS9018 (Corning Inc., US) 96-well plate. Fifty microlitres of the medium were then dispensed into each of the other wells, after which a two-fold serial dilution was carried out. Bacterial suspensions were prepared by touching 3-4 distinct colonies with a sterile cotton swab and dispersing them in PBS. This was then standardised to a 0.5 MacFarland standard with a spectrophotometer (OD600nm = 0.08-0.10). The standardised bacterial suspension (50 µl) was then inoculated into triplicate wells of each concentration except for the sterile control well (without tEDTA or bacteria). Growth control wells with bacteria (without tEDTA) and with no bacteria were also set up. The 96-well plates were sealed with Parafilm™ (Bemis) and incubated at 37°C for 18-24 h. The lowest concentration well with no observable growth was taken as the MIC.

### Biofilm eradication assay in the synthetic collagen wound biofilm model

A synthetic soft-tissue wound model was used as previously described by Werthén et al. (2010). Synthetic wounds were prepared using a mixture of 20% rat tail collagen type 1 (10µg/mL), 60% SWF, 10% of 0.1% acetic acid and 10% of 10mM sodium hydroxide. The mixture was placed on ice and mixed slowly to avoid bubbles. Following this, 200 µL of the mixture was added to wells in a 48-well plate (Greiner bio-one – 677 180). Synthetic wounds were sterilised under short-wave UV-light for 10 minutes before being incubated at 37°C for 1 h to allow the collagen matrix to polymerise. A few colonies of *P. aeruginosa* PA14 from an overnight LB plate were inoculated into 5 ml SWF and incubated with shaking at 37°C for 6 h. This culture was diluted in SWF to obtain an OD_600_ of 0.08-0.1. Fifty microlitres of standardised culture was added to the collagen wound bed and incubated at 37°C for 24 h. After 24 h, 100 µL of tEDTA at 0.5% or 1%, or sterile water (untreated control) was added to the synthetic wounds and incubated at 37°C for 24 h. Following this, 300 µL of collagenase type 1 (0.5 mg/mL) was added to each wound and incubated for 1 h at 37°C to dissolve the collagen matrix. Each well was then mixed thoroughly by pipetting, serially diluted and plated on LB agar plates that were incubated overnight at 37°C. Bacterial colony counts were used to calculate CFU per wound.

### Biofilm eradication assay in the *ex vivo* pig lung model

The *ex vivo* pig lung (EVPL) model has previously been described (Harrison & Diggle, 2016; Harrington et al., 2020; Harrington et al., 2021). Briefly, porcine lungs were obtained from a local butcher (Taylors Butchers, Earlsdon). The bronchioles were dissected and washed in 1:1 RPMI:DMEM with 20 µg/ml ampicillin. The bronchioles were dissected into 5mm strips and washed again in 1:1 RPMI:DMEM. Following this, they were dissected into 5mm squares and washed for a third time in 1:1 RPMI:DMEM and then in synthetic cystic fibrosis sputum media (SCFM1) prepared as described previously (Palmer et al, 2007), but without glucose. Following these washes, bronchiole tissue was sterilised under short-wave UV light in SCFM1 for ten minutes. The tissue sections were placed individually into the wells of a flat, clear 24-well plate (Corning) containing 400 μL of 0.8% w/v agarose in SCFM1. Each piece of tissue was inoculated with PA14 from an overnight LB plate by using a sterile 29G hypodermic needle lightly touch a colony, then to lightly pierce the tissue ∼10 times. After inoculation, 500µl of SCFM1 was added to each well and the plate was sealed with a Breathe-Easier® membrane (Diversified Biotech) and incubated at 37°C for 48 hours. Following incubation, bronchiole tissue with associated biofilm was briefly washed in 500 μL of phosphate-buffered saline (PBS) to remove loosely-adhering planktonic bacteria and placed into a fresh 48-well plate, containing 1%, 2% or 3% tEDTA in SCFM1, or SCFM1 only (untreated control). The plate was sealed with a Breathe-Easier® membrane and incubated at 37°C for 24 hours. Following treatment, the tissue and associated biofilm was briefly washed in 500 μL of PBS and placed into sterile homogenisation tubes each containing 18 2.38mm metal beads (FisherBrand) and 1ml PBS. The tissue was homogenised in a FastPrep-24 5 G (MP Biomedicals) for 40 s at 4 ms^-1^. The homogenate was then serially diluted and plated onto to LB agar, which was incubated at 37°C overnight to determine the CFU per lung piece.

### Biofilm eradication assay in the *in vitro* endotracheal tube model

The *in vitro* endotracheal tube (IVETT) model has been previously described (Walsh et al., 2024a). Endotracheal tubes (ETTS; siliconized PVC, cuffed, 8 mm; IMS Euro) were prepared by cutting a 1cm ring from the tube and chopping this radially into six equal parts under aseptic conditions. Cut sections were sterilised under short-wave UV light for 10 minutes and covered with fetal bovine serum (FBS; Gibco) in a petri dish. This was sealed with Parafilm™ (Bemis) and left at 4°C overnight for serum coating. Serum-coated ETT sections were added individually to the wells of 24-well plates (Corning Costar) using sterile forceps and allowed to dry for 30 minutes. An overnight culture of *P. aeruginosa* PA14 in LB agar was standardised in SVAM (Walsh et al., 2024b) to 0.1 OD_600_, and 0.5 mL was added to each tube section. The plates were then incubated at 37°C, 5% CO_2_ for 24 h, and the biofilm-coated tubes were transferred into fresh SVAM medium for another 24 hours. Tubes were transferred to fresh SVAM containing tEDTA at a concentration of 2%, 3%, or 4%, or to fresh SVAM only (no treatment control) and incubated at 37°C, 5% CO_2_ for 24 hours. Subsequently, the tubes were transferred to 500ml PBS, sonicated at 50 Hz (Grant XUBA1 sonicating water bath) for 15 minutes and agitated further to ensure complete removal of biofilm from the ETT. Serial dilution was performed before plating out on LB agar to determine the colony-forming units per ETT section.

## Conflict of Interest

The authors declare that there is no conflict of interest

## Ethical consideration

No ethical approval was required for this study. Pig lungs were sourced post-slaughter from a commercial abattoir and are thus not relevant material under the Animals (Scientific Procedures) Act 1986.

## Author contribution

Author contributions following CRedit Taxonomy:

Conceptualisation: OO

Investigation: OO, CC, AK, NR, RG

Formal analysis: OO

Methodology: SH.

Supervision: OO, FH.

Writing – original draft: OO.

Writing – review & editing: OO, CC, AK, NR, RG, FH.

## Acknowledgement

OO and AK are funded by the Biotechnology and Biological Sciences Research Council (BBSRC) PhD studentship through the Midlands Integrative Biosciences Training Partnership scheme. OO is also funded by the University of Warwick Institute of Advanced Studies (IAS) Early Career Research Fellowship. CC and NR are funded by the Medical Research Council (MRC) Doctoral Training Partnership. RG is funded by the Engineering and Physical Sciences Research Council (EPSRC) and Tata Steel, UK.

## Notes

### Competing Interest Statement

The authors have declared no competing interest.

